# Basal Internalization and Subcellular Localization of XCR1

**DOI:** 10.64898/2026.06.25.734240

**Authors:** Qiao Li, Fabian Pfersdorf, Fernando Salgado-Polo, Martin Gustavsson

## Abstract

Chemokines orchestrate immune cell trafficking through receptor-mediated signaling and are implicated in inflammatory, autoimmune, and neuropathic disorders. The XCL1–XCR1 axis is of particular interest because XCR1 is selectively expressed on mature conventional type 1 dendritic cells (cDC1s), where it supports communication with activated CD8+ T cells and NK cells and promotes antigen cross-presentation. This selectivity has made XCR1 an attractive target for dendritic cell–based cancer vaccines, while emerging evidence also links XCL1–XCR1 signaling to neuroinflammation and pain.

Despite its therapeutic potential, the mechanisms governing XCR1 activation and trafficking remain understudied. Here, we characterize XCR1 expression, membrane trafficking, and basal internalization to define mechanisms that may influence therapeutic targeting. We show that XCR1 undergoes constitutive internalization through a β-arrestin-independent but adaptor protein 2 (AP2)-dependent pathway, distinguishing it from other chemokine receptors with constitutive endocytosis. Furthermore, we identify specific sequence motifs critical for its subcellular localization and intracellular trafficking. These findings provide new mechanistic insights into XCR1 regulation and may inform the development of targeted therapeutics and antigen-delivery strategies in cancer and inflammation.

## Introduction and Background

Chemokines are key regulators of immune functions and are involved in several proinflammatory and autoimmune diseases. Through the activation of chemokine receptors on the cell surface, chemokines stimulate intracellular signaling pathways, ultimately driving directed cell migration in the direction of chemokine gradients. Chemokines are classified based on the pattern of Cys residues in their N-terminal region, with CC, CXC, CX3C and XC chemokines interacting with CC-CXC- CX3C- and XC receptors, respectively.

Activation of the receptor XCR1 by the XC chemokines XCL1 and XCL2 triggers Gi-coupled intracellular pathways, β-arrestin recruitment and internalization of the receptor [1-3]. The XCL1-XCR1 axis plays a central role in communication between CD8^+^T cells, NK cells, and mature conventional type 1 dendritic cells (cDC1s) [4]. XCR1 is specifically expressed in cDC1s, distinguishing them from precursor cell populations [5]. By secreting XCL1, activated CD8^+^T cells and NK cells recruit mature cDC1 to the inflammation site, promoting antigen cross-presentation [4]. A previous study showed that targeting XCR1 with XCL1 fused to Influenza hemagglutinin (HA) induced stronger CD8+ T cell responses than targeting the more characterized receptors DEC205 and CLEC9A [6]. This suggest that targeting XCR1 on cDC1 may represent a promising, highly selective strategy to deliver antigens in DC-targeted cancer vaccines.

The XCL1-XCR1 axis has also been recently implicated in the nervous system, where it plays a role in the development of neuroinflammation and neuropathic pain [7, 8]. Upon traumatic brain injury in mice, XCL1 levels rise significantly [9], which in turn activate XCR1-expressing dendritic cells, thereby secreting proinflammatory cytokines [8]. Moreover, exogenous administration of XCL1 in the cerebrospinal fluid induces potent and long-lasting pain in naïve mice [10]. Accordingly, neutralization of XCL1 in a diabetic model has been shown to relieve pain, indicating that XCL1-XCR1 signaling mediates neuropathic pain [11].

Although XCR1 has potential as a drug target in several pathophysiological contexts, the mechanisms involved in its activation and intracellular trafficking require further understanding. A recent high-resolution structure identified key interactions with XCL1 that can be used in the production of antagonists [3]. However, XCR1 has been previously reported to internalize constitutively [2], which may limit its availability for drugs on the cell surface. Here, we characterized the expression, membrane trafficking and basal internalization of XCR1. We show that XCR1 internalizes by a β-arrestin-independent and adaptor protein 2 (AP2)-dependent mechanism, different from other constitutively internalizing chemokine receptors. In addition, we identified key sequence motifs involved in intracellular trafficking of the receptor. These findings provide new insights into the function and the subcellular localization of XCR1, which will prove invaluable to inform future therapeutic targeting.

## Materials and Methods

The study was conducted in accordance with the Basic & Clinical Pharmacology & Toxicology policy for experimental and clinical studies [12].

### Cell culture

Parental HEK293A cells, U-87 MG, Chinese Hamster Ovary (CHO) cells, COS-7 cells, or β-arrestin1/2 knockout HEK293A cells were cultured in high-glucose Dulbecco’s Modified Eagle Medium (DMEM) supplemented with 10% fetal bovine serum (FBS) and 1% penicillin/streptomycin at 37 °C and 5% CO_2_ conditions in a humidified incubator.

### Receptor mutagenesis

The cDNA encoding human N-terminally FLAG-SNAP-tagged CXCR4 [13], ACKR3 [13], XCR1 (a kind gift from Mette Rosenkilde, University of Copenhagen) and the mutants thereof were expressed in the pcDNA5.1 vector. For luminescence-based detection, a pcDNA3.1 vector containing the Rluc8 gene was fused at the C-terminus of CXCR4, ACKR3, XCR1 or the mutants thereof using sticky end ligation. Human dynamin 1 K44A (Dyn K44A) was expressed by the pRK5 vector. XCR1 mutants were generated by mutation to the desired amino acid using the Stratagene QuikChange protocol. Mem-citrine was obtained from Jonathan Javitch (Columbia University, NY, USA), and venus-tagged PTP1b, giantin, Rab5, Rab7, Rab11 and MOA were provided by Nevin Lambert (Augusta University, GA, USA). All constructs were verified by bidirectional sequencing.

### BRET-based subcellular localization assay

The BRET ratio between donor-tagged receptors and acceptor-tagged subcellular markers was used to determine the subcellular localization [14] of CXCR4, ACKR3 and XCR1 and the mutants thereof. Fluorescently tagged markers were used to identify different compartments: mem-citrine [15] for the cell membrane, venus-PTP1b [14] for the endoplasmic reticulum, venus-giantin [14] for the Golgi apparatus, venus-Rab5a [16] for early endosome, venus-Rab7a [17] for late endosome, venus-Rab11a [17] for recycling endosome, and venus-MOA [14] for mitochondria.

For this assay, HEK293A cells (30,000 cells per well) were seeded in poly-D-lysine-coated white 96-well plates and co-transfected with 2.5 ng C-terminally Rluc8-tagged receptor construct, 30 ng venus-tagged compartmental protein, and 17.5 ng pcDNA3.1 (for a total amount of 50 ng) using 1 μg/μL polyethyleneimine (PEI) (Polysciences, cat#23966) at a

DNA:PEI ratio of 1:2 (w/v). After 48 hours, the cells were washed twice with PBS and incubated in BRET buffer (PBS supplemented with 0.1% glucose) for 45 minutes at 37 °C. Subsequently, the cells were incubated with BRET substrate (Coelenterazine H, Nanolight technology, cat#301–10) for 8 minutes at a final concentration of 5 μM, after which cells were treated with or without the ligand, and the plate was then measured at 37 °C using an EnVision Plate Reader (Revvity, Inc.)

The subcellular localization of CXCR4, ACKR3 and XCR1 was determined by measuring the BRET ratio between receptor-Rluc8 and each venus-tagged subcellular marker:

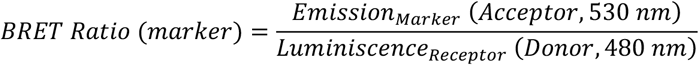

The mitochondrial signal was used as a background control to calculate the fold change for subcellular localization at each compartment:

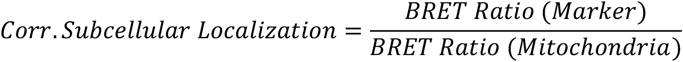

Statistical analysis was performed on the corrected values using ordinary one- or two-way ANOVA with Tukey’s multiple comparisons test.

### DERET-based internalization assay

The internalization assay was performed using diffusion-enhanced resonance energy transfer (DERET), which measures the energy transfer between a SNAP-tagged receptor labelled with a terbium fluorophore probe (SNAP-Lumin4-Tb; Revvity, Inc., cat#SSNPTBD) and a soluble acceptor, Fluorescein-O^’^-acetic acid (Fluorescein; Merck, cat#88596), in the extracellular milieu [18].

For this, HEK293A cells, U-87 MG, CHO cells or COS-7 cells (20,000 cells per well) were seeded in poly-D-lysine-coated white 384-well plates and co-transfected with 0.25-10 ng N-terminally FLAG-SNAP-tagged receptor and 30-39.75 ng pcDNA3.1 (total 40 ng DNA per well) using Lipofectamine™ 2000 at a DNA:Lipofectamine™ ratio of 1:2 (w/v). For experiments with Dyn K44A, 1-10 ng of pcDNA3.1 was replaced by 1-10 ng of Dyn K44A. After 24 hours, the culture medium was removed, and cells were incubated with 100 nM SNAP-Lumi4-Tb at 4 ºC for 1 hour in Opti-MEM™ I Reduced Serum Medium. Subsequently, the cells were washed six times with internalization buffer containing Hanks’ Balanced Salt Solution (HBSS), 1 mM CaCl_2_, 1 mM MgCl_2_, 20 mM HEPES buffer, 0.1% (v/v) Bovine Serum Albumin (BSA), pH 7.4. When indicated, 50 µM Barbadin (MedChemExpress, Cat#HY-119706) or 0.5% DMSO vehicle was added to the wells.

To quantify cell surface expression, internalization buffer was added to the wells, and the signal from SNAP-Lumin4-Tb was measured at 620 nm. To quantify the internalization ratio, fluorescein was added to a final concentration of 25 μM, and the plate was measured at 37 °C for at least 1 h using an EnVision Plate Reader (Revvity, Inc.). The internalization ratio was calculated as follows:

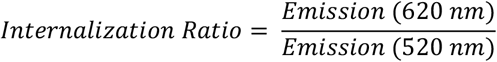

The area under the curve (AUC) for receptor internalization was calculated using GraphPad Prism 10. To correct for the background, the AUC from the group transfected with 40 ng SNAP-pcDNA5.1 was subtracted. Statistical comparisons between the treated and untreated groups were performed using an unpaired t-test. For comparisons involving three or more groups, one-way ANOVA was applied using Tukey’s multiple comparisons post-hoc analysis.

## Results

### XCR1 is a constitutively internalizing receptor with low surface expression

To investigate the surface expression and internalization of XCR1, we transfected HEK293A cells with FLAG- and SNAP-tagged XCR1 or equal amounts of the well-characterized chemokine receptors CXCR4 and ACKR3. CXCR4 is a canonical (G protein-coupled) receptor with low basal internalization and receptors localized mainly at the cell membrane, while ACKR3 is an atypical (not G protein-coupled) chemokine receptor characterized by high basal internalization and a significant portion of receptors localized intracellularly [19, 20]. Labeling of the SNAP-tagged receptors with membrane-impermeable SNAP-Lumi4-Tb showed that XCR1 was expressed at the cell surface, but at levels 35- and 6-fold lower than CXCR4 and ACKR3, respectively (**Fig 1a**).

**Figure 1.**
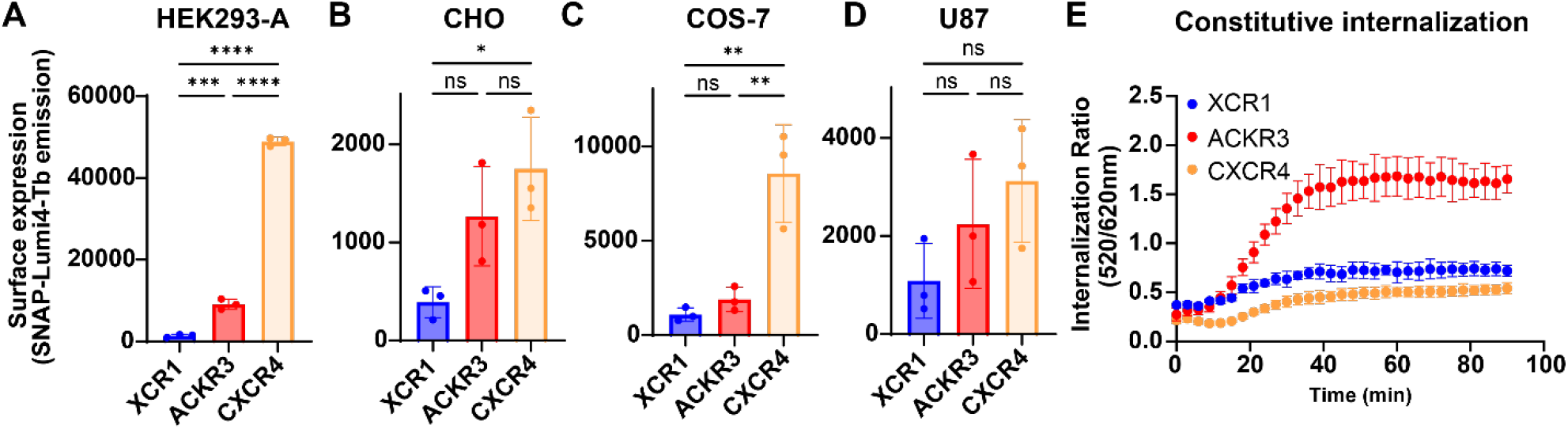
XCR1, CXCR4 and ACKR3 surface expression and basal internalization. (A) Surface expression of Lumi4-Tb-labeled SNAP-tagged receptors in HEK293A cells detected by fluorescence emission at 520 nm. (B-D) Receptor surface expression in CHO (C), COS-7 (D) and U87 (E) cells detected as described in (A). (E) Basal receptor internalization in HEK293A cells detected as the increased ratio of Lumi4-TB emission (520 nm) to fluorescein emission (620 nm) after labeling at 4 °C and transfer to 37 °C. All data show the average of three independent experiments measured in triplicates with error bars corresponding to standard deviation. Statistical analysis was performed with ordinary one-way ANOVA with Tukey’s multiple comparisons test. Asterisks indicate statistically significant differences (ns= p > 0.05, *p < 0.05, **p < 0.01, ***p < 0.001, ****p < 0.0001).

HEK293 cells are a standard cell line for studies of G protein-coupled receptor (GPCR) trafficking and signaling but protein expression can be cell type-dependent and differ between cell lines. To test if the low surface expression of XCR1 is unique to HEK293A cells, we next transfected three other cell lines: Chinese Hamster Ovary (CHO), COS-7 and U87 cells, with SNAP-tagged XCR1, CXCR4 and ACKR3 and quantified receptor surface expression using SNAP-Lumi4-Tb labeling. For all three cell lines, surface labeling of the three receptors is generally lower than in HEK293A cells (**Fig 1b-d**). XCR1 surface expression is significantly lower than CXCR4 in both CHO and COS-7 cells and the general trend, with XCR1 surface expression being low relative to the other receptors, appears to be conserved in all cell lines. This suggests that the low surface expression is a conserved feature of XCR1 and independent of the cell line used.

To assess the basal internalization of XCR1, we next employed a diffusion-enhanced resonance energy transfer (DERET)-based assay, which detects internalization of SNAP- Lumi4-Tb labeled receptors [18]. Time-resolved internalization in HEK293A cells showed that XCR1 internalized more than CXCR4 but less than ACKR3 (**Fig 1e**), indicating that XCR1 internalizes constitutively, but at a lower level than the highly internalizing ACKR3.

Intracellular localization of chemokine receptors may result from an inability to traffic to the cell membrane or from a high basal internalization rate. For ACKR3, its high constitutive internalization causes a significant portion of receptors to be localized intracellularly. Consequently, inhibiting ACKR3 internalization with a dominant negative variant of the small GTPase dynamin (Dyn K44A) prevents basal internalization and leads to increased receptor localization at the cell membrane [13]. To determine whether this also applies to XCR1 we repeated the surface labeling experiments in the presence of increasing amounts of Dyn K44A. Unlike ACKR3, this did not increase surface expression of XCR1 (**Fig S1**), suggesting that the low cell surface expression of XCR1 stems from an inability to reach the plasma membrane.

### XCR1 is retained in the endoplasmic reticulum

To further investigate the expression of XCR1 we next transfected HEK293A cells with receptors (XCR1, ACKR3 and CXCR4) fused to Renilla Luciferase 8 (Rluc8) at their C-termini. Quantification of the luciferase signals showed that despite its low surface expression, total expression of XCR1 was higher in the cell than both ACKR3 and CXCR4 (**Fig 2a**), which further supports the role of membrane trafficking in its low surface expression.

**Figure 2.**
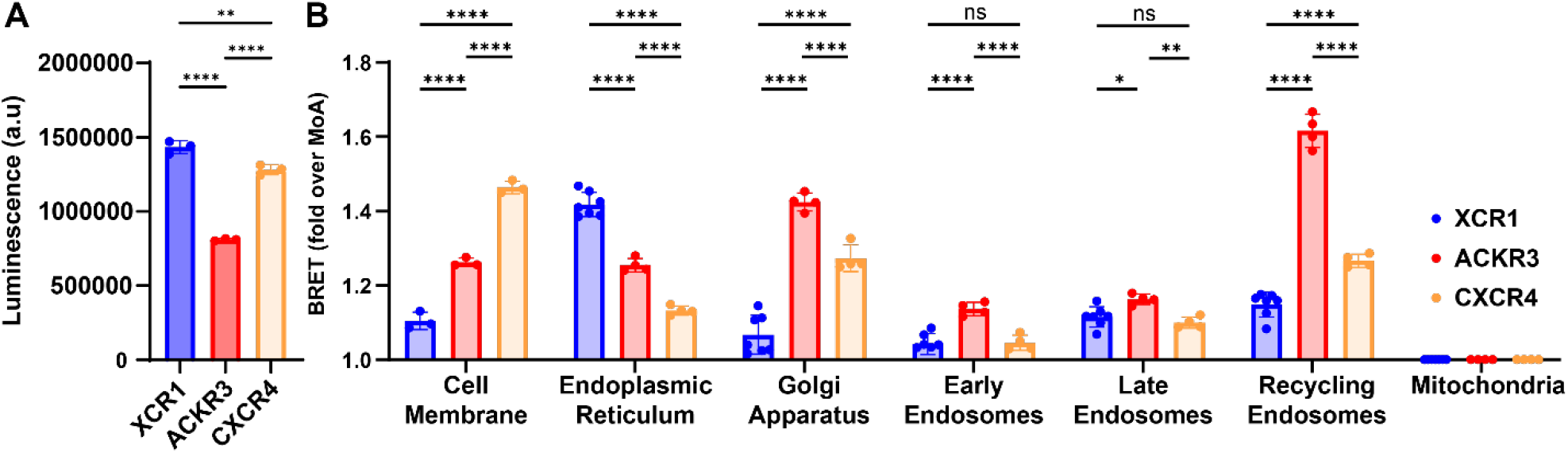
Subcellular localization of receptors. (A) Total luminescence of over-expressed XCR1, ACKR3 and CXCR4 fused to Rluc8 at their C-termini detected by Rluc8 luminescence in HEK293A cells. (B) Localization of receptors in subcellular departments detected by BRET from Rluc8-tagged receptors to fluorescently tagged compartmental markers mem-citrine (cell membrane), venus-PTP1b (ER), venus-giantin (Golgi), venus-Rab5a (early endosomes), venus-Rab 7a (late endosomes) and venus-Rab11a (recycling endosomes) and venus-MOA (mitochondria). All BRET values were normalized to the mitochondrial marker venus-MOA to account for non-specific differences in BRET values. All data are presented as average +/-SD of three independent experiments collected in triplicates. Statistical analysis was performed with ordinary one-way ANOVA (A) or ordinary two-way ANOVA (B) with Tukey’s multiple comparisons test. Asterisks indicate statistically significant differences (ns= p > 0.05, *p < 0.05, **p < 0.01, ****p < 0.0001).

To locate XCR1 and to compare its subcellular localization to the other receptors, we next co-transfected the Rluc8-labelled chemokine receptors with Venus-tagged makers for different intracellular compartments [14-17] and measured BRET to quantify receptor localization in different subcellular compartments (**Fig 2b**). In agreement with the SNAP surface labeling (**Fig 1a**), BRET to the plasma membrane marker Mem-citrine showed that XCR1 is significantly less localized at the cell membrane than both ACKR3 and CXCR4. In contrast, XCR1 had a significantly larger portion of receptors localized to the endoplasmic reticulum (ER) as determined from BRET to the ER marker Ptp1b. In addition, ACKR3 and CXCR4 had significantly higher BRET values with the Golgi apparatus marker giantin. Markers for early (Rab5), late (Rab7) and recycling (Rab11) endosomes show that XCR1 is detected in all three endosomal compartments, but with a different pattern than ACKR3, which had a lower presence in late endosomes and higher in recycling endosomes, and CXCR4, which was more present in recycling endosomes. Taken together, these results suggest that XCR1 is present in multiple intracellular compartments but a large part of translated XCR1 is retained in the ER and not trafficked to the Golgi apparatus for further transport to the plasma membrane, leading to low receptor expression at the plasma membrane.

### Mutation of unique sequence motifs affect XCR1 trafficking and subcellular localization

We next analyzed the sequence of XCR1 to identify unique motifs, which could explain its retention in the ER. The D^3.49^R^3.50^Y^3.51^ (numbers in superscript corresponding to the Ballesteros Weinstein numbering scheme) motif, located at the intracellular end of transmembrane helix 3 (TM3), is highly conserved among Class A GPCRs [21]. In XCR1, the Asp residue of this motif is replaced by His, making it the only chemokine receptor with a H^3.49^R^3.50^Y^3.51^ motif. To address the role of this variation in receptor trafficking, we generated a His-to-Asp acid substitution (H126D) turning the HRY motif of XCR1 into DRY (**Fig 3a**). Compared to wild-type XCR1, the XCR1-H126D mutant had a slightly lower total expression (**Fig 3b**) and a markedly reduced cell surface localization as assessed by SNAP surface labeling (**Fig 3c**) and BRET to the membrane marker Mem-citrine (**Fig 3d**). Analysis of the other compartmental markers showed that XCR1-H126D had higher BRET than XCR1-WT with the ER marker Ptp1b (**Fig 3d**) indicating increased ER retention. In addition, XCR1-H126D was present in late-but not recycling endosomes suggesting that after internalization mutant receptors are degraded rather than recycled to the plasma membrane. Together, this suggests that the HRY motif is crucial for XCR1 folding and/or trafficking from the ER.

**Figure 3.**
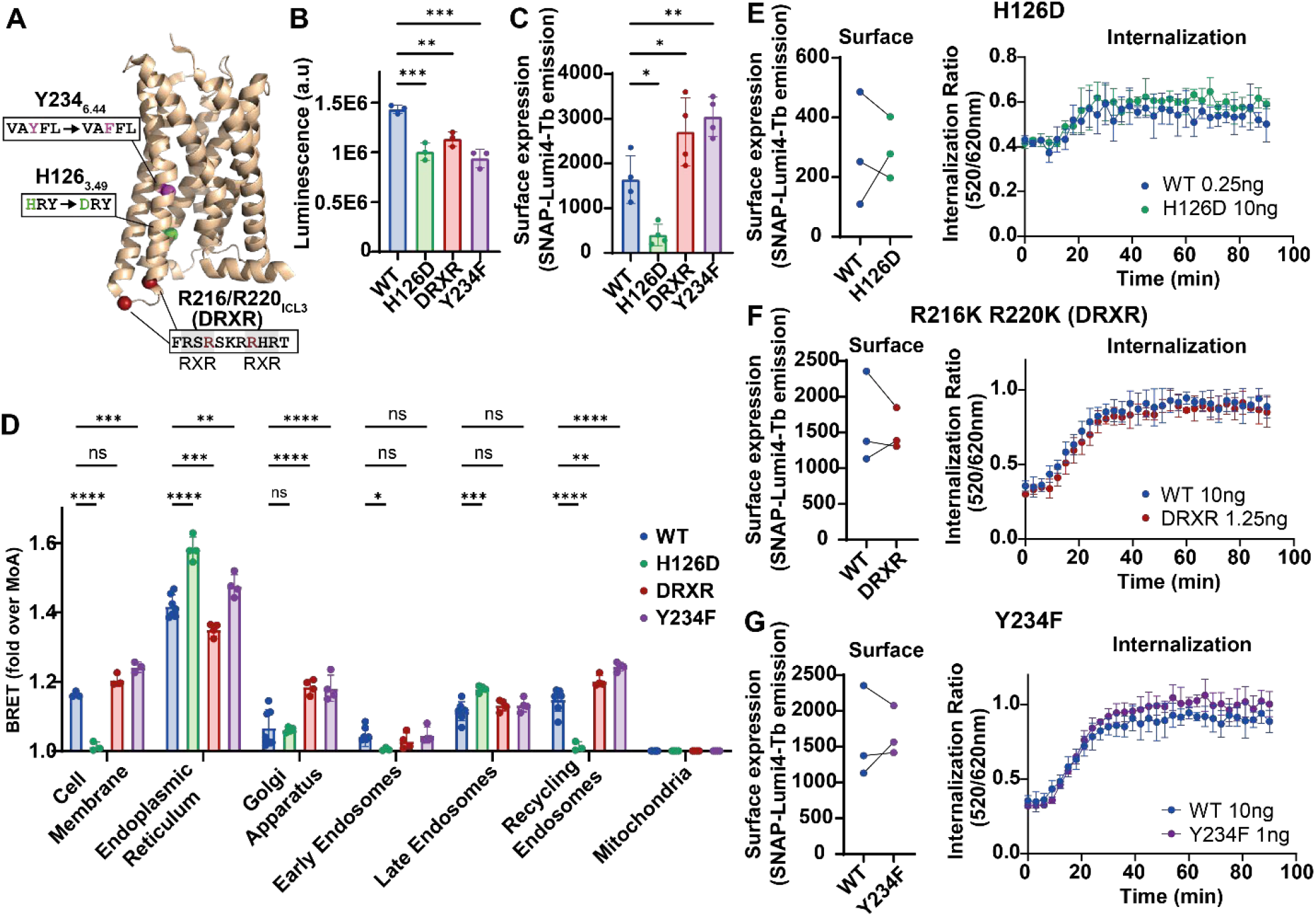
Subcellular localization and internalization of XCR1 mutants. (A) Mutated sites mapped onto the structure of XCR1. (B) Total expression of C-terminally Rluc8-tagged receptors detected by luminescence. (C) Surface expression of receptors detected fluorescence after Lumi4-Tb labeling of SNAP-tagged receptors. (D) Subcellular localization of receptors detected from BRET to compartment-specific Venus-tagged markers. (E-G) Surface expression and internalization of XCR1-WT and XCR1-H126D (E), XCR1-R216K/R220K (DRXR) (F) and XCR1-Y234 (G). XCR1-WT expression in (E) was titrated to match the expression of the mutant receptor and mutant receptor expression in (F) and (G) was titrated to match XCR1-WT with amounts of DNA transfected per well displayed in the figure. Surface expression in (E-G) shown as average of triplicate measurements with lines connecting experiments collected on the same day. All other data are presented as average +/-SD of n=3-4 independent experiments. Statistical analysis was performed with ordinary one-way ANOVA (B-C) or ordinary two-way ANOVA (D) with Tukey’s multiple comparisons test. Asterisks indicate statistically significant differences (ns= p > 0.05, *p < 0.05, **p < 0.01, ***p < 0.001, ****p < 0.0001).

Our sequence analysis also revealed that XCR1 has three Arg-X-Arg (RXR) motifs within the third intracellular loop (ICL3). RXR motifs have been shown to mediate ER retention for other GPCRs [22-24], and we hypothesized that they could contribute to the ER localization of XCR1. To test this, we introduced two Arg-to-Lys substitutions (R216K & R220K) to disrupt both potential ER retention motifs (**Fig 3a**). Although its expression was slightly lower than XCR1-WT (**Fig 3b**), XCR1-R216K/R220K had higher cell surface labeling than the WT receptor (**Fig 3c**). The double mutant also had lower BRET with the ER marker and higher with the Golgi apparatus marker (**Fig 3d**). These findings support our hypothesis and suggest that the RXR motifs contribute to the ER retention of XCR1.

A third unique site is Tyr 234^6.44^ in TM6 of XCR1, which is a Phe residue in most other chemokine receptors and part of the conserved F^6.44^xxCW^6.48^ motif of Class A GPCRs [25] (**Fig 3a**). XCR1-Y234F was recently demonstrated to have an enhanced pharmacological response to XCL1 [3]. The Y234F mutant exhibited a nearly two-fold higher surface expression than that of wild-type XCR1 (**Fig 3c**) despite its overall lower expression (**Fig 3b**). In addition, XCR1-Y234F had a higher amount of BRET than XCR1-WT with markers for multiple intracellular compartments including the ER (**Fig 3d**). This suggests that the increase response of XCR1-Y234F to XCL1 could be explained by an increased surface expression, which in contrast to XCR1-R216K/R220K is not driven by reduced retention in the ER.

Experiments with different transfected amounts of WT-XCR1 showed that receptor internalization depends on surface expression levels (**Fig S2a-c**). To determine how the mutations affected basal internalization of the receptor we therefore titrated the amount of transfected DNA to match surface expression levels for WT and mutant receptors (**Fig 3e-g**). Internalization assays with equal receptor surface expression levels showed that all three mutants had similar levels of basal internalization as WT-XCR1 (**Fig 3e-g**) confirming that the altered subcellular localization is not caused by differences in basal internalization and suggesting that none of the mutants had significant effects on XCR1 internalization. In summary, the three unique motifs influence the overall expression and localization of XCR1 with the RXR motifs promoting the retention of receptors in the ER.

### Basal XCR1 internalization depends on dynamin but not β-arrestins

Previous studies have shown that chemokine receptors differ in their internalization mechanisms. For example, CCR7 depends on β-arrestin for its ligand-induced internalization [26] while the basal as well as ligand-induced internalization of ACKR3 are unaffected in the absence of β-arrestin [27, 28]. We next sought to determine the mechanism of XCR1 constitutive internalization and its potential differences with ACKR3. dynamin is a GTPase required for several internalization pathways, including CME, fast endophilin-mediated endocytosis (FEME), and caveolin-mediated endocytosis [29]. To test its importance for XCR1 internalization we co-transfected SNAP-tagged XCR1 and ACKR3 with Dyn K44A [30]. Receptor internalization was significantly reduced in the presence of Dyn 44A, showing that, as expected, XCR1 internalization is dynamin-dependent (**Fig 4a**) as previously shown for basal internalization of ACKR3 [13] (**Fig 4b**).

**Figure 4.**
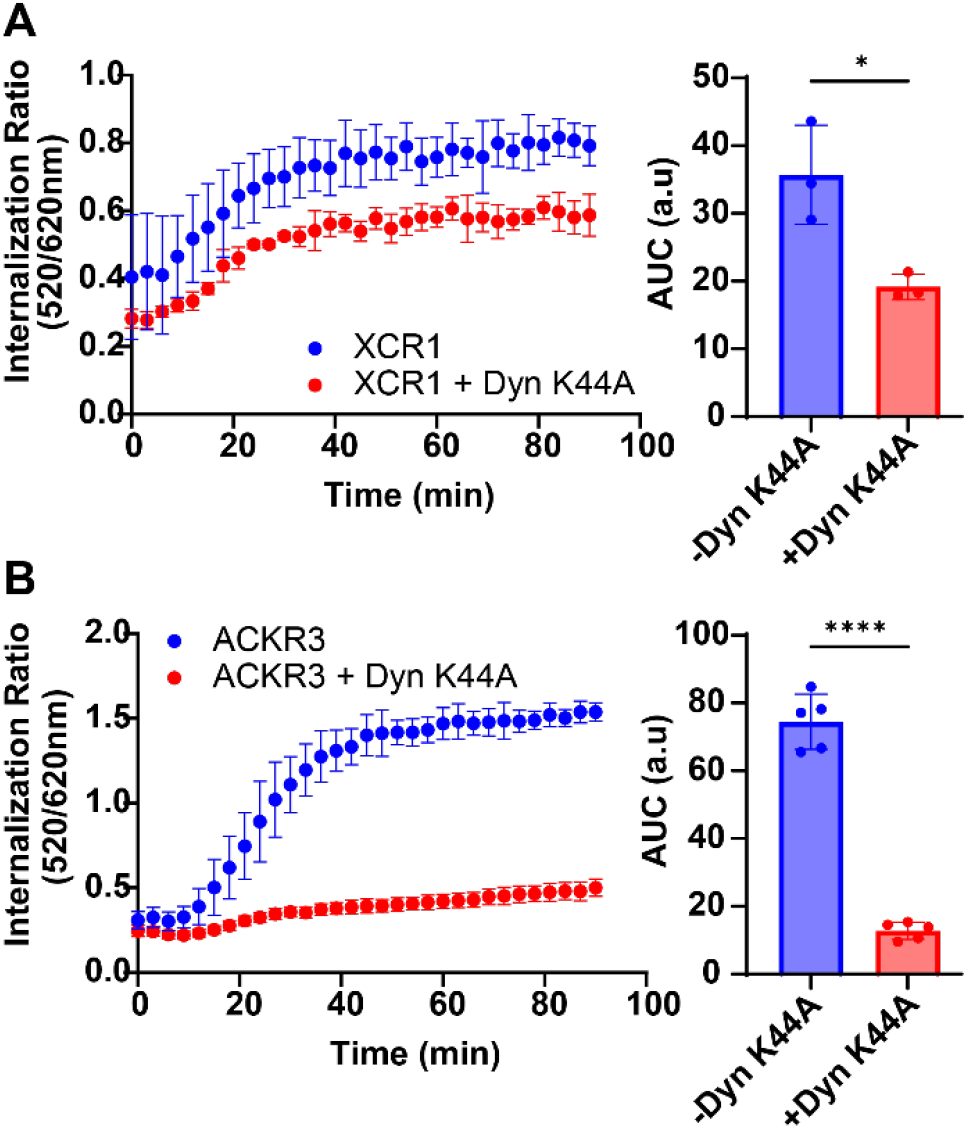
Effect of dominant negative mutant of dynamin on receptor internalization. (A, B) Time-resolved constitutive internalization (left) and area under the curve (AUC, right) determined with or without co-expression of Dyn K44A with SNAP-tagged and Lumi4-Tb-labeled XCR1-WT (A) and ACKR3 (B). Internalization was determined as the increased ratio of Lumi4-Tb emission (520 nm) to fluorescein emission (620 nm) after transfer to 37 °C following labeling at 4 °C. Data correspond to average +/-SD of 3-5 independent experiments collected in triplicates. Statistical comparison was performed using unpaired t-tests *p < 0.05, ****p < 0.0001).

XCR1 is a Gi-coupled receptor, which recruits β-arrestin upon stimulation with XCL1. As β-arrestin mediates internalization of several other GPCRs, we next tested the dependence of XCR1 internalization on β-arrestin using crispr-edited cells lacking both β-arrestin1 and β-arrestin2 (βarr-KO) [31]. Comparison with WT cells showed that XCR1 surface expression and internalization were unchanged in βarr-KO cells (**Fig 5a**), suggesting that XCR1 internalizes independent of arrestin. Similarly, and as previously reported, constitutive ACKR3 internalization was not significantly affected by the absence of arrestins [28] (**Fig 5b**). In contrast, internalization of ACKR5, a chemokine receptor with a high amount of basal internalization, was, as previously suggested [32, 33], strongly dependent on β-arrestin for its basal internalization (**Fig 5c**). Consequently, ACKR5 had a significantly higher surface expression in βarr-KO cells driven by the lower basal internalization (**Fig 5c**). Taken together, this shows that XCR1 internalizes independent of arrestin but that this independence is not conserved across the family of chemokine receptors (**Fig 5d**).

**Figure 5.**
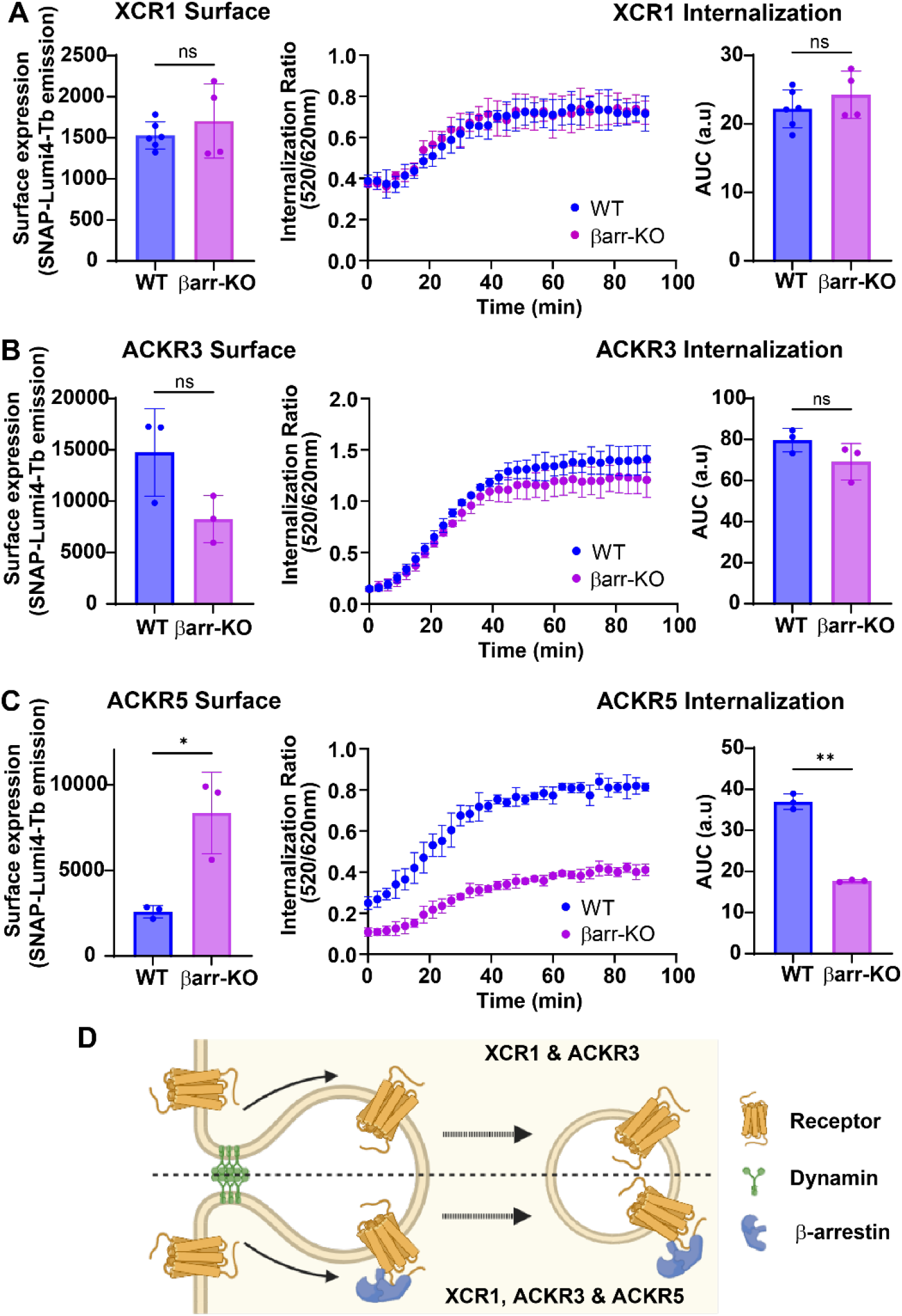
Dependence of XCR1, ACKR3 and ACKR5 surface expression and internalization on β-arrestins. (A-C) Surface expression (left), constitutive internalization (middle) and AUC (right) in WT or βarr-KO HEK293A cells of SNAP-tagged and Lumi4-Tb-labeled XCR1-WT (A), ACKR3 (B) and ACKR5 (C). Surface expression was detected as Lumi4-Tb emission and internalization as the increased ratio of Lumi4-Tb emission (520 nm) to fluorescein emission (620 nm) after transfer to 37 °C following labeling at 4 °C. Data correspond to average +/-SD of three independent experiments collected in triplicates. Statistical comparison was performed using unpaired t-tests (ns= p > 0.05, *p < 0.05, **p < 0.01). (D) XCR1 and ACKR3 internalize independently of β-arrestins, ACKR5 does not. Created with BioRender.com.

### XCR1 internalization is clathrin-mediated

To further pinpoint the mechanism of XCR1 internalization we next the effect of Barbadin, which binds to AP2, a key scaffolding protein in clathrin-mediated internalization [34]. Although Barbadin blocks the interaction between AP2 and arrestins and is generally considered an inhibitor of β-arrestin-dependent internalization, some GPCRs depend on direct interactions with AP2 for their internalization [35, 36]. Surprisingly, Barbadin inhibited constitutive internalization of XCR1 (**Fig 6a**), showing a similar effect to inhibiting all clathrin-mediated endocytosis by Dyn-K44A. In contrast, ACKR3 internalization was only partially inhibited by Barbadin (**Fig 6b**), again showing a difference in internalization mechanisms between the two receptors and excluding that the effects on XCR1 are due to non-specific effects on cellular internalization.

**Figure 6.**
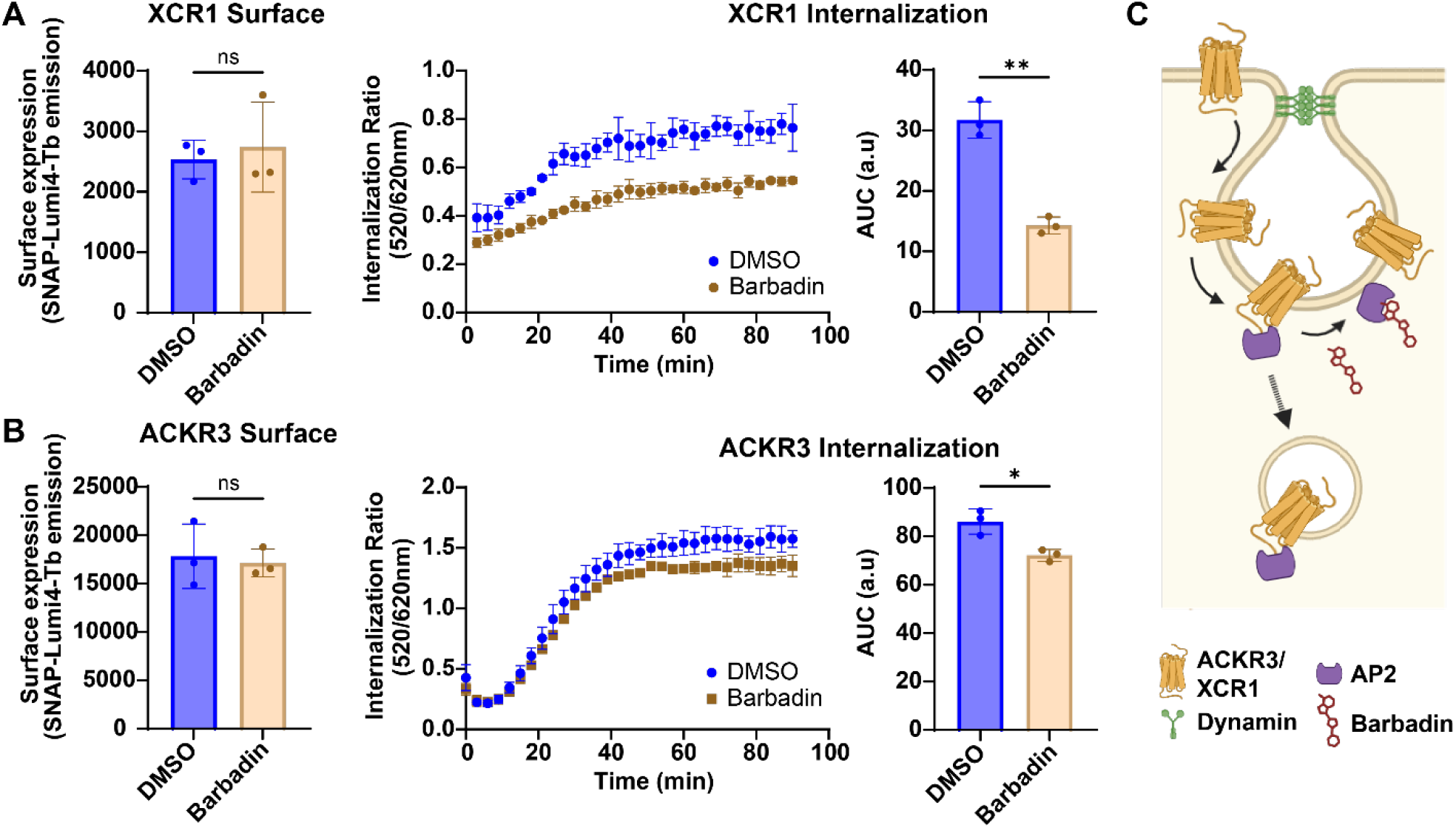
Effects of Barbadin on XCR1 and ACKR3 internalization. (A,B) Surface expression (left), time-resolved constitutive internalization (middle) and AUC (right) after treatment with vehicle (DMSO) or Barbadin of XCR1-WT-(A) and ACKR3-expressing (B) HEK293A cells. Surface expression was detected as Lumi4-Tb emission and internalization as the increased ratio of Lumi4-Tb emission (520 nm) to fluorescein emission (620 nm). Data correspond to average +/-SD of three independent experiments collected in triplicates. Statistical comparison was performed using unpaired t-tests (ns= p > 0.05, *p < 0.05, **p < 0.01). (C) Barbadin binds to AP2 and inhibits basal internalization of XCR1. Created with BioRender.com.

Together, experiments in βarr-KO cells and in the presence of Barbadin show that XCR1 internalizes with a clathrin-dependent mechanism independent of arrestin scaffolding (**Fig 6c**). Instead, XCR1 depends on interactions with AP2, either direct or indirect through an unknown scaffolding protein. This mechanism is different than ACKR3 and ACKR5, two of the other chemokine receptors with significant amount of constitutive internalization. On a chemokine receptor family-level these findings show that the mechanism of basal internalization is receptor-independent and depend on interactions with different scaffold proteins.

## Discussion

XCR1 is unique as the only chemokine receptor binding XCL1 chemokines. Here, we show that XCR1 also has distinct trafficking and internalization properties and depends on unique sequence motifs, further differentiating it from other chemokine receptors. Although most class A GPCRs have a conserved DRY motif in TM3, the presence of His^3.49^ makes XCR1 one of the few receptors with a HRY motif. Mutation of the His^3.49^ residue abolished surface expression with mutant receptors mainly detected in the ER and late endosomes. In inactive GPCR conformations, Asp^3.49^ typically forms a salt bridge with Arg^3.50^, which is broken upon receptor activation allowing Arg^3.50^ to interact with G proteins [37]. Although the positively charged side chain of His^3.49^ should repel Arg^3.50^, our results suggest that XCR1 has adapted to coordinate its (H/D)RY motif, where His^3.49^ is a crucial residue for receptor stability and function.

In addition, mutation of two RXR motifs in the ICL3 of XCR1 increased receptor surface expression. Similar observations have previously been reported for other GPCRs, including PAR4, which is retained in the ER through interactions with COPI-coated vesicles [22]. XCR1 is the only chemokine receptor containing two such motifs in its ICL3, while other chemokine receptors have single RXR motifs in their intracellular loops. However, more studies are needed to determine the mechanism of XCR1 ER retention and the potential role of RXR motifs regulating the trafficking of other chemokine receptors.

It has been previously shown that mutation of Phe^6.44^, part of the conserved F^6.44^xxCW^6.48^ motif, increased pharmacological response to XCL1 [3]. Our results showed that the stronger response of XCR1-Y234F to XCL1 correlates with an increase in the surface expression of XCR1, Furthermore, this exemplifies how characterization of subcellular localization of mutants is a powerful complement to signaling assays to provide a complete mechanistic understanding of receptor function.

Because of its specific expression on subsets of DCs, XCR1 has been proposed as a promising entry port for antigen delivery to cross-presenting DCs [38]. Thus, XCR1 could be exploited in the development of cancer vaccines, making it a potential target in immunotherapy. Similar approaches have previously utilized the virally expressed chemokine receptors US28 and ORF74 to deliver chemokine-fused toxins and selectively kill cells expressing the receptors [39, 40]. US28 and ORF74 are constitutively internalizing receptors, making them well suited for intracellular toxin delivery and opening the possibility to utilize both chemokine agonist and antagonists for efficient cell killing. Here, we show that XCR1, like US28 and ORF74, is a constitutively internalizing receptor, consistent with a previous report for the rat isoform of XCR1, potentially making it more favorable as a vehicle for antigen delivery.

Previous studies have also shown that the efficiency of receptor-mediated antigen cross-presentation depends on trafficking to different endosomal compartments, with a preference for delivery to early endosomes compared to late and recycling endosomes [41]. Our results showed that XCR1 can be detected in all endosomal compartments, although at relatively low levels in early endosomes. Furthermore, we showed that XCR1 internalizes with a clathrin-mediated endocytosis. Clathrin-coated vesicles can deliver cargo up to 100-200 nm in size [42] and are an efficient route for antigen processing [43]. Together, these findings support the potential role of XCR1 in cancer vaccine development.

Basal internalization of XCR1 and, as previously reported, ACKR3 [27, 28] occur independently of arrestins. In contrast, the constitutive internalization of ACKR5, recently introduced as the latest member of the chemokine receptor family, was strongly arrestin-dependent. This is in agreement with previous observations using knockdown or knockout cell lines for arrestin expression [32, 33]. Although many GPCRs depend on arrestins for ligand-induced internalization, a recent study suggested that basal internalization is generally arrestin-independent [44]. In fact, although a majority of 60 tested GPCRs depended on arrestins for ligand-dependent internalization, none of them depended on arrestin to internalize in the absence of ligand. This makes ACKR5 unique among the GPCRs tested for their arrestin-dependence. However, most studies of GPCRs have focused on ligand-induced signaling and trafficking, and future studies may identify other receptors with similar properties. Overall, our results show that multiple mechanisms can drive constitutive internalization of chemokine receptors.

Although independent of arrestins, XCR1 constitutive internalization was inhibited by Barbadin, suggesting that AP2 is crucial for its internalization. Previous studies have detected multiple GPCRs where direct interactions of AP2 with the C-terminal tail of the receptor contribute to its endocytosis. For example, ligand-dependent internalization of α1b-AR depends on AP2 interactions with a stretch of eight consecutive Arg residues [45], constitutive internalization of PAR1 is mediated by AP2 association with a YXXL motif [35], ligand-induced internalization of GLP-1 receptor is partially dependent on AP2 interacting with a GRK-phosphorylated stretch of Ser residues [44] and mutation of a LLKIL sequence in CXCR2 reduced AP2 interaction and ligand-induced internalization [36]. Our results suggest that XCR1 may interact directly with AP2; however, XCR1 does not contain any of the previously established AP2-binding motifs in its C-terminal domain. Thus, more studies are needed to establish the molecular mechanisms of its AP2 recruitment, including whether the interactions are direct or part of a larger protein complex. In addition, although several chemokine receptors internalize independently of arrestins, the potential role of AP2 in basal- and ligand-induced internalization of chemokine receptors has yet to be explored.

Our characterization of XCR1 was mainly done in HEK293A cells and we also confirmed that XCR1 had a low surface expression also in CHO, COS-7 and U87 cell lines. The use of these immortalized cell lines was crucial to enable the various biosensors and molecular tools used in our study. However, future studies in primary cells naturally expressing XCR1 will be crucial for a complete understanding of the surface expression and trafficking of XCR1.

In summary, we have identified and tested sequence motifs and mechanisms to provide new insights into the basal trafficking and internalization of XCR1. Our results establish unique features of the receptor and could pave the way for targeting of XCR1 by cancer vaccines or other therapeutic approaches.

## Acknowledgements

We thank Rajesh Regmi for expert technical assistance. This work was supported by Carlsberg Foundation grant CF19-0320 (M.G), Villum Fonden grant 00025326 (M.G), Independent Research Fund Denmark grant 3103-00230B (M.G) and Danish Cancer Society grant #R389-A22939 (F.S.-P)

