## Supplemental figures 1 and 2 for "Basal Internalization and Subcellular Localization of XCR1"

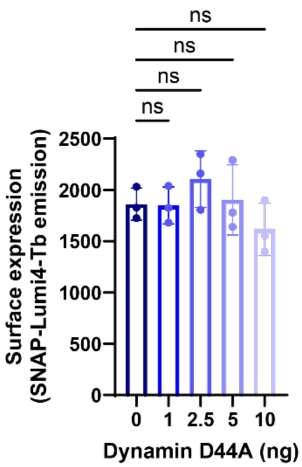


**Figure S1**. Surface expression of XCR1 after co-expression of increasing amounts of Dyn K44A. Statistical analysis was performed with ordinary one-way ANOVA with Tukey's multiple comparisons test. (ns $p > 0.05$)

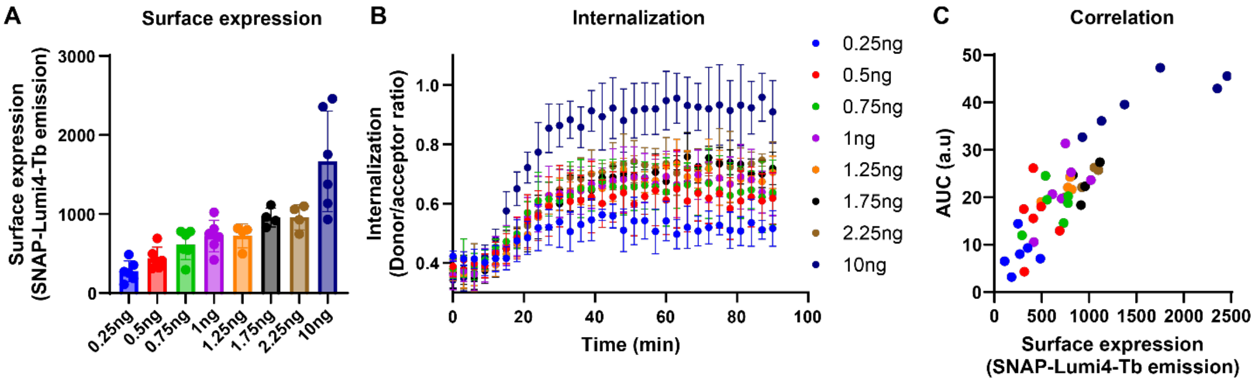


**Figure S2**. (A) Surface expression after transfection with different amounts of XCR1 DNA. (B) internalization with different amounts of transfected XCR1 DNA. Data correspond to average +/- SD of 4-6 independent experiments collected in triplicates. (C) Correlation between surface expression and area-under the curve of internalization for each replicate displayed in (A).
